# Antibody Complementarity Determining Region Design Using High-Capacity Machine Learning

**DOI:** 10.1101/682880

**Authors:** Ge Liu, Haoyang Zeng, Jonas Mueller, Brandon Carter, Ziheng Wang, Jonas Schilz, Geraldine Horny, Michael E. Birnbaum, Stefan Ewert, David K. Gifford

## Abstract

The precise targeting of antibodies and other protein therapeutics is required for their proper function and the elimination of deleterious off-target effects. Often the molecular structure of a therapeutic target is unknown and randomized methods are used to design antibodies without a model that relates antibody sequence to desired properties. Here we present a machine learning method that can design human Immunoglobulin G (IgG) antibodies with target affinities that are superior to candidates from phage display panning experiments within a limited design budget. We also demonstrate that machine learning can improve target-specificity by the modular composition of models from different experimental campaigns, enabling a new integrative approach to improving target specificity. Our results suggest a new path for the discovery of therapeutic molecules by demonstrating that predictive and differentiable models of antibody binding can be learned from high-throughput experimental data without the need for target structural data.

**Significance:** Antibody based therapeutics must meet both affinity and specificity metrics, and existing *in vitro* methods for meeting these metrics are based upon randomization and empirical testing. We demonstrate that with sufficient target-specific training data machine learning can suggest novel antibody variable domain sequences that are superior to those observed during training. Our machine learning method does not require any target structural information. We further show that data from disparate antibody campaigns can be combined by machine learning to improve antibody specificity.

## Main Text

The identification of human antibodies and receptors with high affinity and specificity to human disease associated targets is a key challenge in producing effective human therapeutics. At present antibody sequences are discovered *in vivo* using animal immunization or by *in vitro* affinity selection of candidates from large synthetic libraries of antibody sequences ^1–5^. These methods both have the advantage that they do not require the structure of a target to be known for antibodies to be discovered. However, they are empirical and do not produce a model of sequence space that admits computational optimization and specificity analysis without conducting additional experiments such as counter panning.

Most computational methods for antibody design optimization assume that the structure of a target is known, however target structure is unknown is many cases^6^. Other approaches seek to optimize antibody properties primarily focused on predicting the structural conformation of the CDR-H3 loop, which to date remains a difficult unsolved challenge ^7,8^. Many such approaches are based on calculations of binding free energies, where the multitude of possible expressions have been found to be of highly variable quality for predicting actual affinities in experiments ^9,10^. More recently, neural network methods have been successfully applied to predict which region of the CDR will be in contact with an antigen (without reliance on structure prediction), but this work does not provide insights on how to obtain improved affinity ^11^. Existing models of antibody binding do not produce gradients for sequence optimization that directly relate antibody sequence to a desired objective.

Here we describe the application of high-capacity machine learning to design antibody sequences, rather than merely predicting their properties. Unlike other computational approaches, the molecular structure of an antigen does not need to be known, permitting our method to be applied to any target used in affinity selections. Three recent developments have enabled this approach. First, high-throughput sequencing^12^ has made it possible to develop large training data sets that describe the relationship between antibody sequence and antibody affinity enrichment with respect to a target of interest. Second, high capacity machine learning approaches such as neural networks and ensembles^13^ enable large antibody sequence-enrichment data sets to be summarized as predictive and differentiable machine learning models that can be used to generate optimal sequences more efficiently compared to random optimization approaches. Third, new array-based DNA oligo synthesis technology^14^ permits the experimental testing of millions of machine-learning proposed designs, both to find optimal candidates and to probe sequence space to build better models for sequence synthesis.

Our hypothesis is that with sufficient sequence-only training data, high-capacity machine learning can sufficiently model the biophysics of antibody-target interactions to generalize from these observations to produce novel improved antibody sequences. High-capacity machine learning methods have a large number of parameters that permit them to model complex relationships with rich expressive power ^15^. The high-capacity models we use are differentiable, meaning that we can compute the derivative of an objective function with respect to an input sequence, allowing for direct gradient-based optimization of the input with respect to the objective. We tested our hypothesis by generating training data that relates Fab fragments with varying CDR-H3 sequences to their enrichment in phage display panning experiments with four target antigens and one mock control. Chosen from publicly available molecules to perform panning experiment against, our four targets consisted of the antibodies ranibizumab (Fab fragment binding to vascular endothelial growth factor A), bevacizumab (monoclonal antibody binding to vascular endothelial growth factor A), etanercept (a fusion protein that fuses the TNF receptor 2 to the Fc and hinge region of a IgG1 heavy chain), and trastuzumab (monoclonal antibody binding to human epidermal growth factor receptor 2). We chose antibodies as targets because they include well characterized common and unique regions that allow us to characterize the ability of machine learning to learn region specific binders. In addition, anti-antibody molecules are important as therapeutic reversal agents^16^. Our mock control consisted of panning with no antigen and thus measures CDR-H3 sequence specific bias for display on phage and phage propagation.

We first sought to determine if we could increase antibody target-specificity by using machine learning to reject antibody sequences that bind to other undesired targets. We found that we could use neural networks trained on other Fc-region containing targets including trastuzumab and etanercept to predict antibodies that were enriched for bevacizumab but not for the other Fc-region containing targets. This result demonstrates that models learned from multiple targets can be combined and transferred to a new target to select antibodies with desired properties.

We next explored if we could model antibody affinity with an ensemble of neural networks and efficiently optimize it with gradient based optimization (Ens-Grad). We also examined alternative machine learning approaches based upon a variational auto-encoder (VAE) and genetic algorithms (GA-CNN, GA-KNN). We tested these diverse methods to examine how best to design new antibody CDR-H3 sequences that bind to ranibizumab based upon high-throughput training data. We discovered that the neural network ensemble approach produced accurate predictive estimates of affinity enrichment for previously unseen sequences, and that with a small design budget of 5,467 sequences gradient based optimization of ensemble-predictions produced antibody sequences that were superior to all of the sequences present in our training data. We interpreted our machine learning models by computing the minimal sets of specific CDR-H3 amino acids required for binding^17^. Our results demonstrate that machine learning can produce useful models of antibody affinity that can create novel sequences with desired properties and increase target specificity.

## Results

### Machine learning predicts panning enrichment

To select Fab sequences against ranibizumab we conducted three rounds of phage display panning (Fig. 1A). In parallel, we conducted so-called mock panning in which we infected and propagated phages without contact to an antigen. The synthetic input library for panning contained a fixed framework with fixed CDR sequences except for CDR-H3. CDR-H3 was randomized with ~10^10^ different sequences that ranged in length from 10 to 18 amino acids. The phage output of each panning round (R1, R2, and R3) was analyzed by Next Generation Sequencing at the CDR-H3 locus and the frequency of each CDR-H3 sequence was computed as its fractional representation in the round. We obtained 572,647 unique CDR-H3 sequences for ranibizumab in round 1, 297,290 in round 2 and 171,568 in round 3. We define *enrichment* as the log_10_ of the round-to-round ratio of sequence frequencies, where positive enrichment indicates affinity selection of a sequence from round to subsequent round. We use R2-to-R3 enrichment for training as it had a higher signal-to-noise ratio. Both positive and negative enrichment of a subset of CDR-H3 sequences was observed for ranibizumab (Fig. 1C and 1D). We observed preferred CDR-H3 lengths for ranibizumab binding (Fig. 1E). Moreover, the CDR-H3 sequences with high enrichment form isolated clusters that correspond to distinct enriched sequence families. We visualize these clusters in 2D using parametric t-SNE dimensionality reduction (Fig 1F, Fig. S1, and Supplementary Materials).

**Fig. 1.**
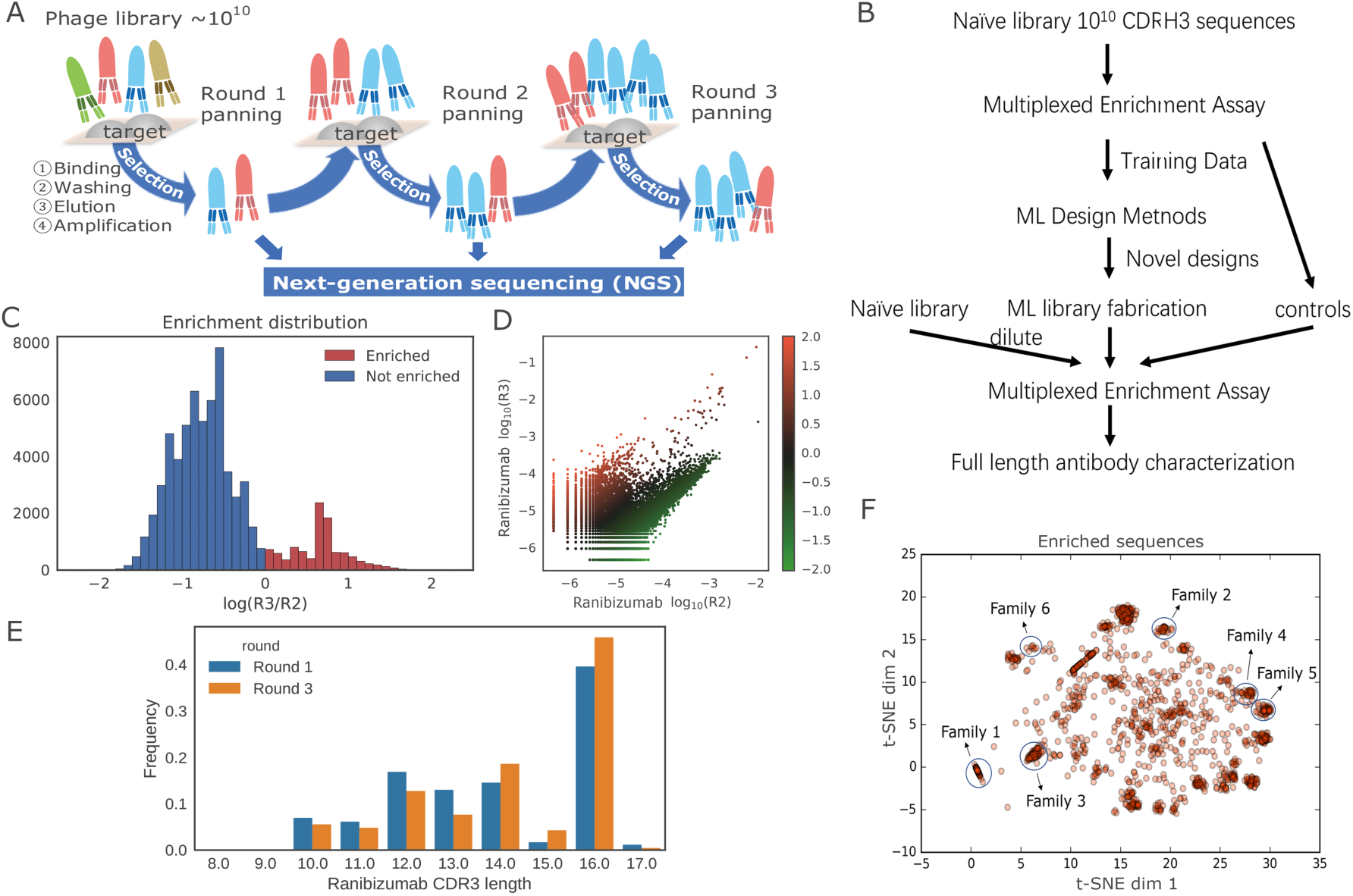
Summary of the training data used in our machine learning framework. (A) Diagram of the training data generation using phage display panning and NGS. Three rounds of panning (R1, R2, R3) were performed. We characterize the frequency of CDR-H3 sequences at the ends of each panning round to compute enrichment. (B) Diagram of the whole workflow. (C) Histogram of R2-to-R3 enrichment for ranibizumab in training data. R2-to-R3 enrichment is defined as log10 of R2-to-R3 frequency ratio. Y-axis denotes the sequence counts in each bin. (D) Scatter plot of log10 sequence frequency in R2 and R3, colored by the R2-to-R3 enrichment value. Each point represents a unique valid sequence in the NGS output, where points above the diagonal have positive R2-to-R3 enrichment and vice versa. (E) Histogram of CDR-H3 sequence length before and after 2 rounds of panning. Y-axis denotes the proportion of reads in each bin. Ranibizumab binders (survived sequences in round 3) tend to exhibit greater CDR-H3 lengths (compared to sequences in round 1). (F) t-SNE visualization of sequences with over 10-fold enrichment between round 2 and round 3. CDR-H3 sequences with high enrichment form isolated clusters. Sequences similar to the ones in Table 1 are labeled by circles. CDR-H3 sequences with high enrichment form isolated clusters.

We trained classification and regression machine learning models on R2-to-R3 enrichment. To train the classification model we assigned binary labels to sequences as either binding or not binding to an antigen based on R2-to-R3 sequence enrichment (Supplementary Materials). We excluded training examples that showed little positive or negative enrichment for classification. To train the regression model we used the real-valued observed R2-to-R3 enrichment. We did two experimental replicates for panning against ranibizumab and the correlation of enrichment between identical sequences that appeared in both replicates was 0.83 (Pearson *r*, Fig. 2A). We trained our models on replicate I and used non-overlapping sequences in replicate II as the test set for ranibizumab. For our Mock control, we randomly held out 20% of the CDR-H3 sequences for testing.

**Fig. 2.**
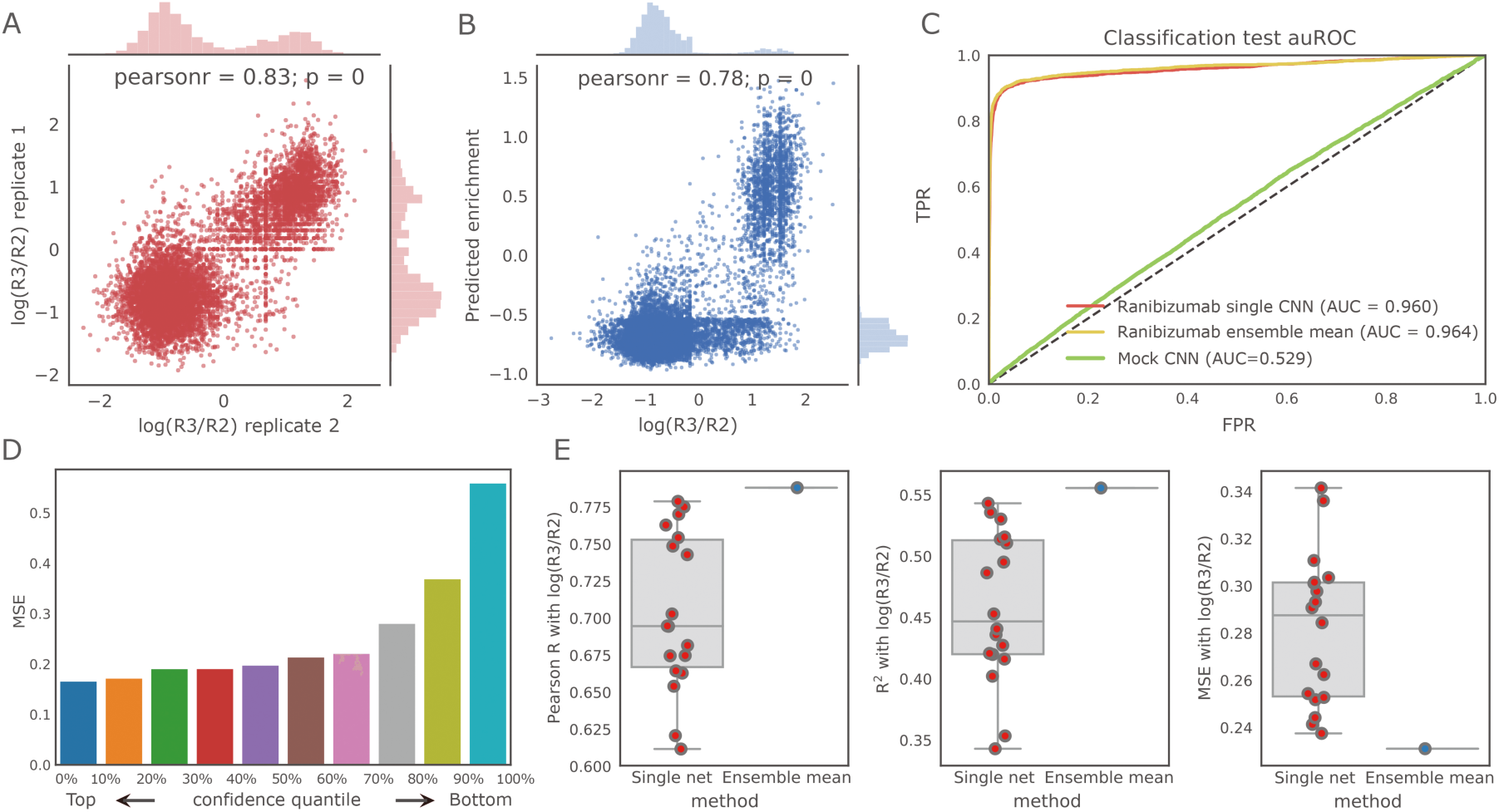
Performance of our neural networks in predicting ranibizumab enrichment. (A) Regression performance for ranibizumab on non-overlapping sequences in replicate II using the best single network (Pearson r = 0.78, p < 1e-16, n = 20,775). (B) Replicate consistency of enrichment for ranibizumab (Pearson r = 0.83, p < 1e-16, n = 9,117); the Pearson r is about the same between replicates as between the neural network predictions and the observed enrichment values, suggesting that our prediction performance is bounded by the replicate consistency. (C) ROC curve of classification task for ranibizumab using the best single network and the ensemble, compared with a mock control. The ensemble method outperforms a single neural network, while the performance in mock is close to random guessing, as expected. (D) Uncertainty estimates of ensemble correlate with prediction accuracy. Sequences assigned higher uncertainty tend to have greater mean square error. (E) Ensembles (blue) produce better estimates of enrichments than any single network (red) in terms of Pearson r, R2, and mean square error. Boxplots show median (the center line in the box), 25th and 75th percentiles (the boundaries of the box), 1.5 interquartile range (the ends of the whiskers), and outliers (points outside of the whiskers). Uncertainty estimates of ensemble correlate with prediction accuracy. Sequences assigned higher uncertainty tend to have greater mean square error.

We found that our classification and regression models were able to accurately predict the R2-to-R3 enrichment of held-out test sequences. Using a convolutional neural network, test set classification performance was 0.960 AUROC (Area Under the Receiver Operating Curve), and test set regression performance was 0.78 (Pearson *r*) (Fig. 2B and 2C). Given there is a 0.83 correlation (Pearson *r*) between experimental replicates of naïve library panning for R2-to-R3 enrichment, the performance of the regression model is bounded by experimental noise (Fig. 2A). Performance on the classification mock control was 0.529 AUROC and mock regression performance was 0.025 (Pearson *r*). These mock results are consistent with our expectation of no predictive ability when attempting to infer phage enrichment in the absence of target binding.

To test the model’s capability to extrapolate beyond best seen sequences, we built an independent held-out validation set consisting of top 4% of the all observed sequences using R2-to-R3 enrichment. We trained a convolutional neural network for regression task on the lower 96% sequences, and compared the predicted scores between the held-out top sequences and top 4% of training sequences. Our model assigned higher scores to the unseen sequences when compared to top seen sequences, indicating that convolutional neural network is able to extrapolate beyond seen examples. We also observed that the network was able to assign higher score to the upper half of the held-out set when compared to the lower half (Fig. S2).

### Ensembles of neural networks improve accuracy and characterize uncertainty

A method that produces novel antibody sequences will naturally need to venture in sequence space outside of observed training data. Thus, when evaluating sequences, it is important to estimate both the expected value of enrichment as well as the model uncertainty of the estimated expectations. We accomplish this with an ensemble of neural network models of diverse architectures that have been trained on different sets of observations. Ensemble learning is a well-established strategy to provide robust uncertainty estimates for the predictions of deep learning methods ^18–20^.

We created an ensemble of neural networks by training 18 different regression models to predict CDR-H3 enrichment using six different architectures and three different training sets for each architecture. The six architectures include five convolutional neural networks and one fully connected network (Table S1, Supplementary Materials).

We first asked whether the uncertainty encoded in the ensemble correlates with prediction accuracy. We quantified the prediction accuracy using mean-square-error (MSE) and estimated ensemble uncertainty by computing the variance of the predictions of the models in the ensemble. With the ensemble trained on panning replicate I, we evaluated the uncertainty and accuracy on sequences only observed in replicate II. We observed a strong positive correlation between the mean-square-error and the variance of the ensemble prediction (Fig. 2D), showing that confident predictions from the ensemble are more accurate.

We hypothesized that the mean of an ensemble of models would result in a more robust estimator of sequence enrichment than using prediction from a single network. Relying on the predictions of a single-model is risky in settings involving noisy measurements and a massive feature space that is only sparsely populated by the training data. We compared the performance of our 18 single models with an ensemble method that averaged the prediction of all 18 models. Evaluating on our held-out data we observed that the ensemble model outperformed all of our 18 models in all metrics considered including Pearson *r*, *R*^2^ and mean-squared error (Fig. 2E).

### Machine learning can eliminate non-specific antibodies that bind to undesired targets

We hypothesized we could create a model to identify antibodies that bind etanercept or trastuzumab, and then use this model to reject anti-bevacizumab antibodies that bind to etanercept, trastuzumab, or the IgG Fc region they share with bevacizumab. We conducted three rounds of phage display panning (Fig. 1A) separately against etanercept, trastuzumab and bevacizumab as targets starting with the same randomized library (Methods). We analyzed the phage output of each panning round (R1, R2, and R3) by sequencing the CDR-H3 locus and computed the frequency and R2-to-R3 enrichment of each CDR-H3 sequence.

To predict antibodies that were enriched for bevacizumab but not target-specific, we trained an ensemble of multi-output neural networks with the six architectures defined in Table S1 which output predicted R2-to-R3 enrichment of both etanercept and trastuzumab (Supplementary Materials). We randomly held out 5% of the training data as test set to evaluate model performance. We found that our regression models were able to accurately predict the R2-to-R3 enrichment of held-out test sequences for both etanercept and trastuzumab. Using the ensemble mean as prediction, test set regression performance was 0.65 (Pearson r) for trastuzumab and 0.64 (Pearson r) for etanercept (Fig. 3A). We calculated the ensemble variance to incorporate model uncertainty, and used the 95% confidence upper bound of the predicted scores for trastuzumab and etanercept to indicate potential non-specific binding.

**Fig. 3.**
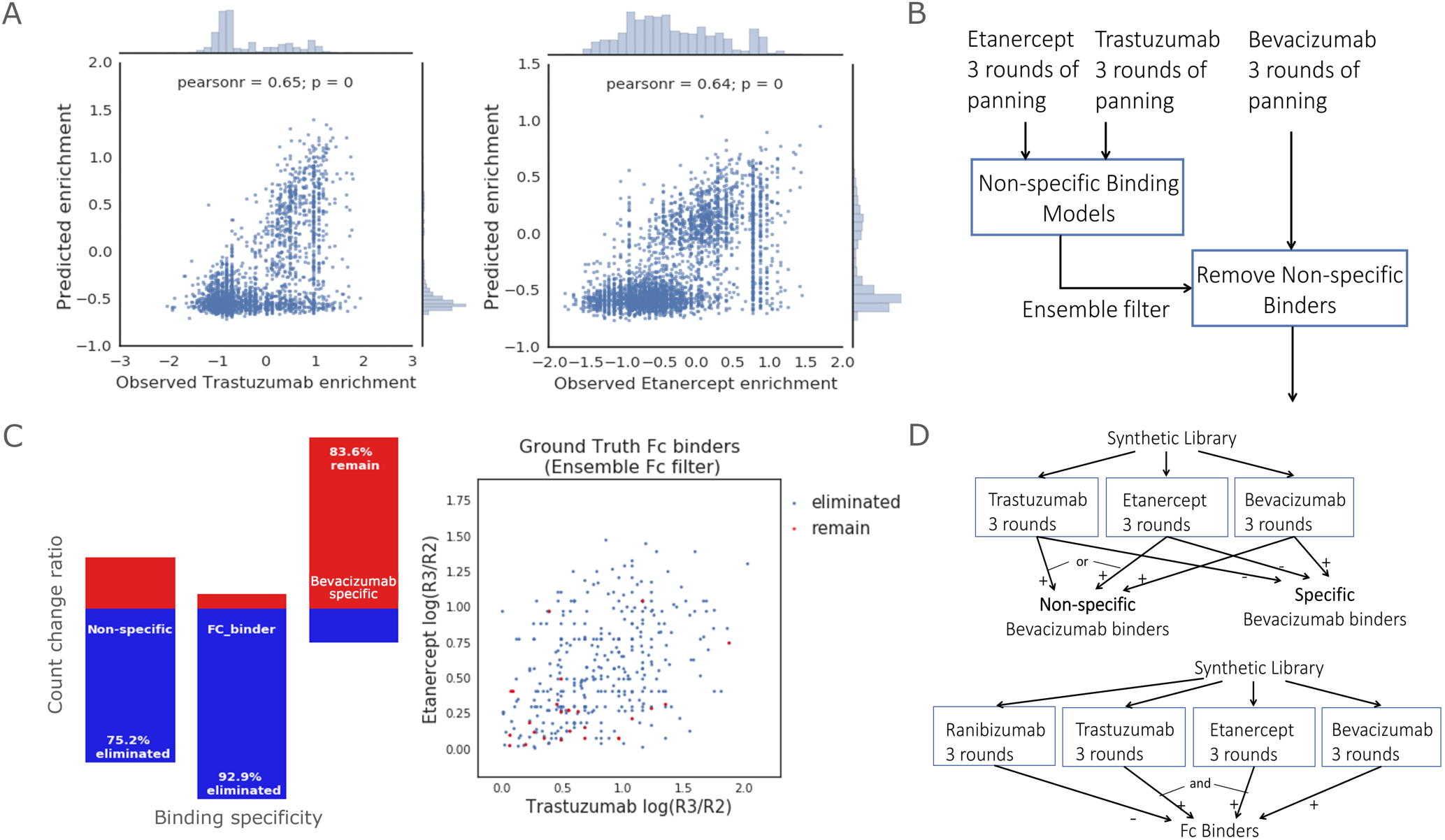
Machine learning can reject undesired antibodies. (A) Regression performance of a multi-output network on a randomly held-out 5% test set with no-missing values for trastuzumab (left, Pearson r = 0.65, p < 1e-16, n = 2,971) and for etanercept (right, Pearson r = 0.64, p < 1e-16, n = 3,487). (B) Diagram of the non-specific binding prediction task. (C) Changes in count of non-specific antibodies, FC-binders and bevacizumab-specific antibodies after filtering antibodies that were enriched for bevacizumab using the specificity prediction ensemble (left). Bevacizumab binding antibodies plotted vs. their etanercept and trastuzumab enrichment, both of which contain an Fc region (right). Blue dots represent ground truth bevacizumab Fc binders that were eliminated and red dots represent Fc binders remained after applying the corresponding filter on all bevacizumab binders. By using the ensemble prediction, we successfully removed 92.9% of the ground truth bevacizumab Fc binders, which is significantly more than random down-sampling of bevacizumab binders (p-value < 1e-16). We found 83.6% of specific bevacizumab binders remained after filtering, showing that the procedure successfully reduces Fc binding without eliminating most non-Fc binders. (D) Definition of non-specific, specific bevacizumab binders and Fc binders using panning experimental data. Positive sign represents positive R2-to-R3 enrichment and negative sign represents non-positive enrichment.

We then applied our non-specific prediction ensemble on the antibodies that were enriched for bevacizumab to examine their binding specificity (Fig. 3B). Our bevacizumab data were not used to train the non-specific prediction model, and we randomly excluded 50% of the bevacizumab binders that also appeared in the etanercept and trastuzumab experiments from the training set and test our non-specificity elimination performance on these held out sequences. For all antibodies that were enriched for bevacizumab, we filtered out potential non-specific antibodies that had a positive confidence upper bound score for either etanercept or trastuzumab. Our non-specific prediction ensemble successfully eliminated 75.2% of general non-specific binding to either trastuzumab or etanercept (Fig. 3C, Table S2, p < 1e-16). We found that the percentage of bevacizumab-specific antibodies was increased by 7.4%, and 83.6% of the bevacizumab-specific antibodies remained after filtering (Fig. 3C), showing that the procedure successfully reduces non-specific binding while retaining most target-specific binding.

We defined 366 sequences as ground truth IgG Fc binders that showed positive R2-to-R3 enrichment in bevacizumab, etanercept and trastuzumab (all Fc containing) but non-positive enrichment in ranibizumab (no Fc region). Our non-specific prediction ensemble successfully eliminated 340 of the 366 Fc binders (92.9%) and the remaining sequences have low R2-to-R3 enrichment (Fig. 3C, p < 1e-16).

### Machine learning designed sequences are better than panning derived sequences

Our next aim was to examine whether computational models are capable of designing previously unseen CDR-H3 sequences with improved affinity for ranibizumab. We employed a gradient-based optimization framework (Ens-Grad) to propose high-affinity sequences from an ensemble of neural networks trained to predict enrichment. For comparison, we also applied alternative computational approaches to propose improved sequences including a variational autoencoder (VAE) and genetic algorithms (GA-KNN, GA-CNN; Supplementary Materials). We selected sequences with either non-negative R2-to-R3 enrichment or larger than 5e-5 frequency in R3 as seed sequences that were used to initialize the optimization for proposing novel sequences. Both the seeds and sequences that the neural networks predicted to be negative were subsequently included as experimental controls to validate our computational methods and to provide baseline performance. Collectively we constructed a library of 104,525 sequences for experimental testing (Methods).

Ens-Grad’s neural network ensemble for optimizing sequences consists of 24 neural networks based on six network architectures and four different training sets for each architecture (Fig. 4A, Table S1). The training data included three different regression training sets that were partially overlapping, and one classification training set. Optimization of an input sequence by a neural network was accomplished by the backpropagation of gradients that updated the input of each stage of the network to increase its output score. This use of backpropagation is in contrast to its usual application of updating network parameters as in this application network parameters are held constant. In each case, the optimization was started at a given seed sequence. During gradient ascent, we relax the constraint that each input to the neural network must be strictly one-hot (a discrete amino acid sequence) and the optimization was instead conducted over a continuous input space using initial step size λ. After every k iterations, the current continuous input representation is projected into a one-hot representation by selecting the amino acid at each position with the maximum value (Fig. 4B). The resulting sequence is then evaluated and compared to the best recorded previously projected sequences from the same seed. If the newly projected sequence fails to improve upon the best score for 10 iterations we terminate the optimization, presuming it has converged to local optimum of the network’s score function in one-hot space.

**Fig. 4.**
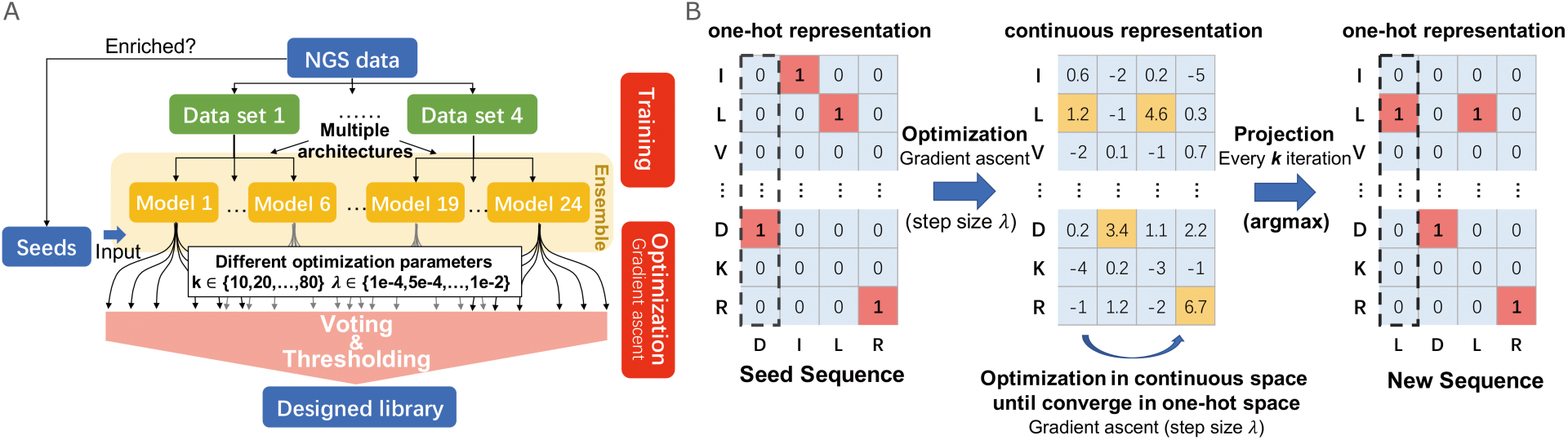
Overview of the Ens-Grad sequence-optimization methodology. (A) Neural network ensemble optimization and thresholding pipeline. (B) Optimization of a single sequence in one-hot space and continuous space using gradient ascent. After every k iterations of gradient ascent in continuous space, we project the current continuous sequence representation back to the nearest one-hot vector by selecting at each sequence position whichever amino acid has received the largest value.

In Ens-Grad, each individual network from the ensemble proposes improved sequences using specified configurations of parameters k and λ during the sequence-optimization with respect to the predictions of this network. All of the proposed sequences are subsequently combined into a candidate pool. We then used the entire ensemble to score each candidate sequence via majority voting and selected the most reliable sequences as our final candidates. Sequences that failed to pass the last filtering stage were also used as control sequences to validate the predictive power of the ensemble (Supplementary Materials, Fig. 4A).

We experimentally tested the synthetic sequences in a three-round phage display panning experiment using both stringent and standard washing conditions. We created DNA oligonucleotides from the sequences generated by Ens-Grad, the alternative computational methods (GA-KNN, GA-CNN, and VAE), the seed sequences, and the negative controls after enforcing several compositional constraints to eliminate chances of deamidation or becoming glycosylated and to ensure the CDR sequences can fold normally (Methods). We cloned them into a phage Fab expression framework with the other CDR sequences fixed to their values during training data generation, and created phage that displayed this library. This phage library was diluted 1:100 into a synthetic library of complexity ~10^10, such that the library complexity was equivalent to the previous experiment and our designed sequences were not underrepresented. We used the 1:100 library in a three-round phage display panning experiment and analyzed the R1-to-R3 enrichment to characterize the binding affinity of the computationally designed sequences when compared with the seed sequences and negative controls that were also present in the oligo derived phage library. We grouped all of the tested sequences, including controls and optimized sequences, by their observed R1-to-R3 enrichment and found that our ensemble of neural networks correctly assigned the highest prediction scores to the top group, whereas sequences that showed negative enrichment were appropriately assigned the lowest scores (Fig. 5A).

**Fig. 5.**
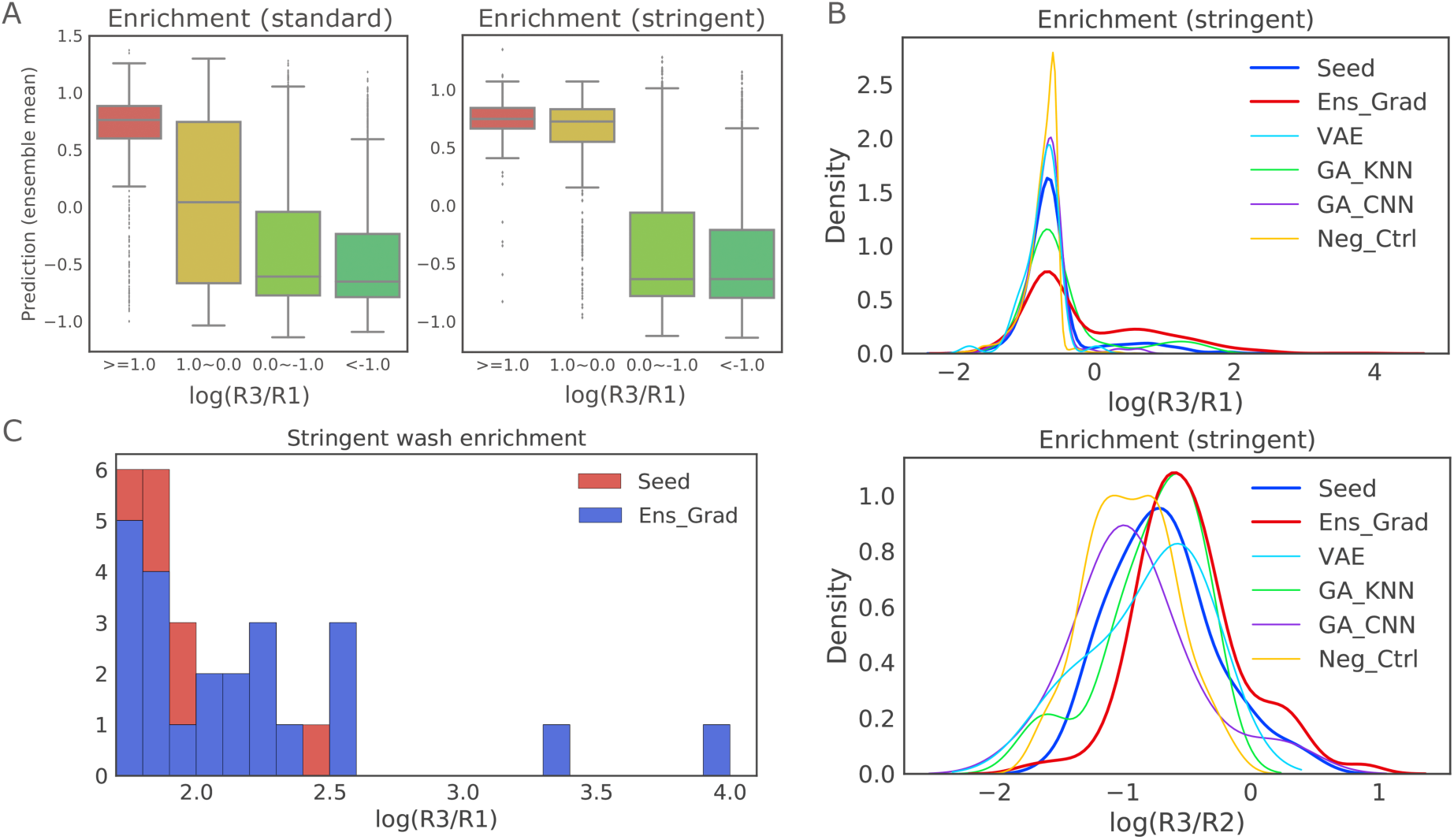
Experimental results of the ML-proposed library diluted 1:100 into our initial synthetic library. (A) Correlation between predicted enrichment using ensemble and actual R1-to-R3 enrichment of all sequences in the synthesized library, including optimal sequences, seed sequences and negative controls. Boxplots show median (the center line in the box), 25th and 75th percentiles (the boundaries of the box), 1.5 interquartile range (the ends of the whiskers), and outliers (points outside of the whiskers). (B) Distribution of R1-to-R3 and R2-to-R3 enrichment of sequences proposed by each of the methods, together with seed sequences and negative controls. Ens-Grad proposed sequences that are better in general by introducing a positive distributional shift compared to seeds, whereas the other methods failed to improve upon the seed sequences. (C) Stacked histogram of the right tail of R1-to-R3 enrichment distribution of each method. Ens-Grad sequences were in higher percentage on the right tail, and the best Ens-Grad sequence outperformed the best seed sequences by 28-fold. None of the sequences designed by other methods attained enrichment higher than 1.5 and thus don’t show up in this
plot.

We compared the enrichment of the computationally-optimized sequences and the seeds in the three-round phage display panning experiment. During analysis we excluded seed sequences that might arise from sequencing noise, and the designed sequences optimized from them (Supplemental Materials). We observed that the 5,467 sequences proposed by ensemble gradient ascent method (Ens-Grad) show a positive distributional shift of R1-to-R3 enrichment compared to seed sequences (two-sided Mann–Whitney U test; for standard washing condition, U = 372780, p = 5.324e-13, and difference in location (Hodges–Lehmann estimate) = 0.097 with 95% confidence interval = [0.079, 0.146]; for stringent washing condition, U = 204640, p = 1.333e-13, difference in location = 0.146 with 95% confidence interval = [0.097, 0.176]) (Fig. 5B). With stringent washing, the mean R1-to-R3 enrichment value of Ens-Grad derived sequences is 0.341 log10 higher than that of the seeds, indicating an improvement equivalent to 62.13% of the standard deviation of seeds R1-to-R3 enrichment (0.549). The Ens-Grad method also produced the sequences with the highest enrichment at the upper tail of the enrichment distribution (Fig. 5C), outperforming all top sequences in seeds by 28-fold (1.450 in log10 fold). Ens-Grad produced sequences with up to 5 changes away from seeds, with Ens-Grad sequences with 2 changes producing sequences with more than a hundredfold R1-to-R3 enrichments (Fig. S3C). We observed negative or insignificant positive distributional shift (two-sided Mann– Whitney U test; p = 0.025 and difference in location = −0.067 for VAE, p = 0.140 and difference in location = −4.619e-5 for GA-CNN; p = 0.776 and difference in location = 2.214e-5 for GA-KNN) of R1-to-R3 enrichment in sequences produced by each of the 3 alternative baseline methods. Similar patterns were observed for the distribution of R2-to-R3 enrichment. The limitation in performance suggests that the genetic algorithm could not adequately explore the optimal sequence space and the latent space in autoencoder based methods could not fully capture the correspondence between valid antibody sequences and their enrichment.

We selected a subset of the top sequences including 12 seeds and 7 machine learning-designed sequences from six different sequence families and synthesized complete IgG molecules of these sequences to evaluate their affinity using an ELISA EC50 assay (Supplementary Materials). The EC50 analysis included the top seed sequences and machine learning-designed sequences within each family, as measured by R1-to-R3 enrichment using both standard washing and stringent washing. Within each CDR-H3 sequence family, we observed that the machine learning-designed sequences had superior or equivalent affinity for ranibizumab than any of the seed sequences, with the best machine learning-designed antibody having an EC50 of 0.29 nM (Table 1, Fig. S3B). We found that a subset of CDR-H3 amino acid positions and values were Sufficient Input Subsets for Ens-Grad to predict enrichment at a threshold of 0.4 (Table 1) ^17^. We also found Ens-Grad preferred to change certain CDR-H3 amino acid positions to optimize enrichment, and the Ens-Grad Sufficient Input Subsets necessary for enrichment diverged from the simple observed frequency of optimized sequences (Supp. Figure S6).

**Table 1.**
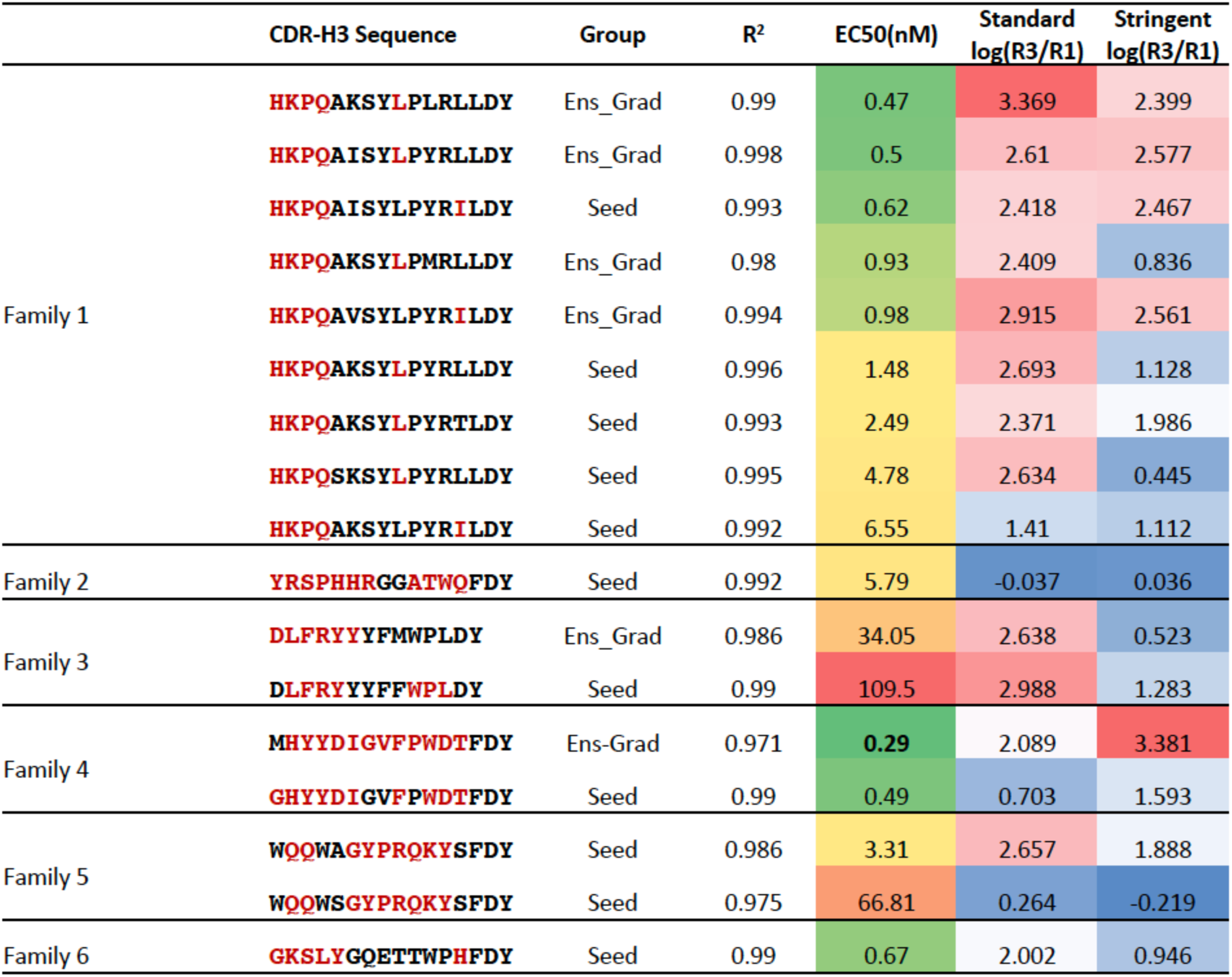
Examples of the top ML-proposed sequences and top seed sequences, along with their enrichment in standard and stringent washing condition, EC50 affinity measurement, and R^2^ fit of EC50 model. We observed higher correlation between phage panning enrichment and EC50 using stringent washing condition. Ens-Grad-proposed sequences outperformed the best seed sequences in each example sequence family in both enrichment and EC50. Colored in red are the positions that, according to our neural network ensemble, contribute the most toward the strong enrichment of these sequences. Each shaded region comprises a Sufficient Input Subset, which alone provides sufficient evidence for the ensemble to justify a highly positive prediction >= 0.4 of enrichment.

## Discussion

We have found that machine learning based methods are an effective way to both model and optimize antibody complementarity-determining region sequences based upon experimental training data. We note that our methods are not intended as an alternative to conventional randomized affinity maturation that can explore tens of millions of alternative candidates. Rather, machine learning provides a modular method to combine models based on data from both primary antibody campaigns and subsequent affinity maturation steps to achieve desired affinity and specify objectives. We observe that within a single antibody campaign machine learning can produce optimized sequences that are on average and at the extremes better than what they observe during training. As DNA sequencing of panning rounds can be performed at a low cost our methods can be readily applied to new antibody campaigns to discover CDR sequences that improve affinity and specificity using the methods we have described.

Among the various learning algorithms applied to generate optimized CDR-H3 sequences, we found that Ens-Grad was able to produce sequences with the highest binding affinity. The factors underlying the superior performance of Ens-Grad include its utilization of neural networks to accurately predict sequence affinity, an ensemble of many networks to ensure robustness in the face of predictive uncertainty, and gradient ascent applied directly over the space of one-hot encoded amino acid sequences rather than more complex optimization strategies. Learning from phage panning experiments that affinity purify a large synthetic library, the Ens-Grad methodology proposed here is able to efficiently generate sequences whose affinity exceeds that of the entire synthetic library comprising the training data.

We demonstrated that we can combine machine learning models to reject antibodies that bind undesired targets, demonstrating that we can improve antibody binding specificity with data from disparate targets. Antibody specificity is a key property for therapeutic safety, and we expect that multi-objective machine learning models will permit the identification of desirable therapeutic candidates. Importantly, compared to experimental procedures such as counter panning which is specific to an undesired target, multi-objective machine learning models can be readily applied to improve antibody specificity by building upon data from past antibody campaigns.

Our results suggest machine learning will provide efficient strategies for exploring promising subsets of sequence space for antibody design. Conventional affinity maturation methods, when sufficiently powered, improve the affinity of CDR-H3 sequences by randomization. We note that Ens-Grad proposed sequences with 2 amino acid changes from panning derived sequences which exhibited 1.899 log10 R1-to-R3 enrichment in stringent washing and 2.888 log10 R1-to-R3 enrichment in standard washing (Fig. S3C). A brute force search of all possible 1 and 2 changes for 6566 seed sequences would require up to 2.193× 10^8^ sequences, while Ens-Grad explored certain of these changes within a design budget of 5,467 sequences. Thus, machine learning offers a powerful tool for focusing design candidates on optimal sequences.

We expect that further experiments that test machine learning designed sequences can add to the corpus of training data, improving predictive models. Bayesian Optimization based acquisition functions can be employed to make informed decisions about where to sample in sequence space to update models to produce better predictions and sequence suggestions. Such suggestions can be used as the seeds for conventional affinity maturation strategies. An attractive property of machine learning based models is that they can be generalized to predict the enrichment of a sequence for more than one target in a multi-task framework. Multi-task frameworks naturally allow for the multi-objective optimization of antibody sequences for both affinity and specificity. The Sufficient Input Subsets of CDR-H3 sequences our model predicts are key for experimental enrichment provide a rich set of hypotheses for future structural studies.

In the presence of adequate training data, we expect that the general optimization framework we outline will be applicable to a wide range of design challenges including the design of DNA aptamers, RNA aptamers, and general protein sequences. We expect that our methods will work unchanged for other CDRs and with data from other affinity platforms such as yeast display^21^. The convergence of high-throughput experimental data from multiplexed experiments, high-capacity machine learning that can process these data, and the direct synthesis of machine learning suggested candidate molecules offers an attractive area for further exploration.

## Acknowledgments

We thank Siddhartha Jain, Nathan Hunt, and Hidde Ploegh for constructive discussions. Funding was provided by NIH grants R01CA218094, and the Novartis Institutes of Biomedical Research;

## Author Contributions

D.K.G., G.L. and H.Z. initiated the study; J.S., G.H., and S.E. performed the phage-display panning and ELISA experiment; the phage-display panning data was processed and analyzed by G.L.; the ELISA experiment data was processed and analyzed by J.S., G.H., and S.E.; D.K.G., G.L., H.Z., J.M., and Z.W. devised the computational models and designed the computationally-optimized sequences; the interpretability analysis was performed by B.C.; the manuscript was prepared by G.L., H.Z., J.M., B.C., and D.K.G. All authors discussed the results and commented on the manuscript;

