## Supplementary Information for "Antibody Complementarity Determining Region Design Using High-Capacity Machine Learning"

### Methods

#### Single framework library generation

A gene fragment encoding the germline framework combination IGHV3-23 and IGKV1-39 was synthesized by Invitrogen's GeneArt service in Fab format and cloned into a phagemid vector serving as the base template. IGHV3-23 and IGKV1-39 were used as they display a favorable framework combination for a phage display library and this framework combination is used in several therapeutic antibodies including trastuzumab and bevacizumab[22]. The phagemid vector consists of Ampicillin resistance, ColE1 origin, M13 origin and a bi-cistronic expression cassette under a lac promotor with OmpA - light chain followed by PhoA–heavy chain – Amber stop – truncated pIII (amino acids 231 – 406). Only CDR-H3 was diversified and primers were designed to incorporate all naturally occurring amino acids excluding cysteine (free cysteines could form disulfide bonds), and asparagine (asparagine in conjunction with certain amino acids could undergo deamidation or become glycosylated) using trinucleotide technology (ELLA Biotech). Lengths between 10 and 16 amino acids and 18 amino acids were allowed, in which the last two amino acids were kept constant with the sequence Asp-Tyr for length 10 to 16 and Asp-Val for length 18. The design of the final two CDR-H3 amino acids reflects human VDJ recombination. Short CDR-H3s more often use J-fragment IGHJ4 with "DY" at the end of CDR-H3 while longer CDR-H3s (here 18 aa) more often use IGHJ6 with "DV" at the end of CDR-H3. Library inserts were generated by PCR using Phusion High Fidelity DNA polymerase (NEB Biolabs). The resulting CDR-H3 library inserts were ligated into the base template, transformed into E.coli TG1 DUO (Lucigen) with a minimal library size of 1E+09 transformants per CDR-H3 length and phages were produced using M13KO7 helper phage (NEB Biolabs) using standard previously described protocols [23].

#### IgG expression

Candidates of interest were expressed in human IgG1 format as previously described [24]. In brief, genes of light and heavy chains were synthesized (GeneArt), separately cloned into pRS5a plasmid and co-transfected using PEI MAX (Polysciences) into CAP-T cell line (CEVEC Pharmaceuticals). Supernatants were purified using protein A affinity chromatography. Purified samples were checked on SDS-PAGE and analytical size exclusion chromatography.

#### Phage Display panning against target molecule

Panning of the library against target etanercept, trastuzumab, bevacizumab, and ranibizumab (in-house expression and purification) was done in solid phase mode [23]. 96-well maxisorb plates (Nunc) were coated with the target using 500 nM in first and second round and 200 nM in the third round and then blocked with PBS/5% (w/v) dried milk protein. 1E12 phages (approximately 50x the library size) were blocked in 0.5x Chemiblocker (Millipore)/2.5% (w/v) dried milk protein containing 0.05% (v/v) Tween-20 and applied on the coated wells. The washing regimen was intensified during the selection rounds with 5 cycles (3x quick and 2x 5 minutes) using PBS with 0.05% Tween 20 followed by 5 cycles (3x quick and 2x 5 minutes) using PBS in the first round, 5 cycles (1x quick and 4x 5 minutes) using PBS with 0.05% Tween 20 followed by 5 cycles (1x quick and 4x 5 minutes) using PBS in the second round and 15 cycles (10x quick and 5x 5 minutes) using PBS with 0.05% Tween 20 followed by 15 cycles (10x quick and 5x 5 minutes) using PBS in the third round. For the stringent washing condition the same number of washing cycles were used but with a buffer containing 0.75M NaCl, 2% Octyl β-D-glucopyranoside and 0.1M Tris, pH 7.4 while the last two steps of each regimen were done with PBS. Phages were eluted with 10mM Glycine/HCl pH 2.0. E.coli TG1F+ were infected with eluted phages which were neutralized beforehand using 1M Tris/HCl, pH8.0. Propagation of phages between rounds was performed using VCSM13 helper phage (Agilent).

**Next Generation Sequencing (NGS) sample preparation**

After each round of phage selection, polyclonal plasmid DNA was prepared using QIAprep Spin Miniprep Kit (Qiagen). In a first PCR the CDR-H3 was amplified over 9 cycles with overhanging primers allowing the annealing of the standard TruSeq primers in a consecutive step. In the second PCR primers with the TruSeq universal forward adapter and one of the 24 TrueSeq index reverse primer were used over 12 cycles. The resulting fragment was purified using a 1.5% agarose gel and the Wizard SV Gel and PCR Clean-up System (Promega). DNA concentration was measured using the Qubit DNA High sensitivity kit (Invitrogen). Samples were analyzed on a MiSeq using MiSeq Reagent Kit v3 (Illumina) with 150 forward cycles or on a HiSeq using HiSeq PE Cluster Kit v4 cBot and HiSeq SBS Kit v4 (Illumina) with 76 forward cycles.

**ELISA**

MaxiSorp black plates (Nunc) were coated with 30nM of ranibizumab at 4 °C overnight. Wells were washed 3 times with TBS containing 0.05% (v/v) Tween-20, blocked with TBS/5% (w/v) dried milk protein for 1 h, washed again and incubated with a serial dilution of purified anti-ranibizumab human IgG1 in TBS containing 0.05% (v/v) Tween-20, 0.5% (w/v) dried milk protein for 1 h. Following four wash cycles, detection was achieved using PO-conjugated Anti-Human IgG Fc (Sigma). The reaction was developed using BM Chemiluminescence ELISA POD Substrate (Roche) and read in a TECAN Genios Pro Reader (TECAN) by luminescence. EC50 and standard variation values were calculated using a 3-parameter logistic regression fit using Prism 7.03 Software (GraphPad).

**Data preparation**

We obtained ~10^7 HiSeq raw reads of length 76 of the nucleotide sequences around CDR-H3 region for ranibizumab from all 3 rounds of panning, 4,493,746 reads from round 1, 3,983,153 reads from round 2 and 3,526,752 reads from round 3. The fixed flanking sequences on the boundary of variable region (12 base pairs on the head, 9 base pairs on the tail) was used as template to locate and segment out the CDR-H3 sequence. We used BLAST [25] for short reads alignment to align the template with each read, allowing maximum of 2 mismatches on each side. We then took the sequence between end position of head template and start position of tail template, and discarded it if the length was not a multiple of three. On average, 58.90% of the raw reads were successfully parsed. The translation to amino acid sequences was done using EMBOSS [26]. The resulting sequences were checked again for abnormal length (smaller than 8 or longer than 20 amino acids) or a stop codon. We obtained 572,647 unique CDR-H3 sequences for ranibizumab in round 1, 297,290 in round 2 and 171,568 in round 3. The same sequencing and preprocessing procedures were used for the other three targets and details can be found in Table S2. In the replicate of the same experiment, we obtained 8,022,860 MiSeq raw reads of length 150, in which over 85% were successfully parsed. In the replicate the number of unique CDR-H3 sequences is 558,400 for round 1, 265,563 for round 2 and 96,912 for round 3. We obtained 1,895,225 MiSeq raw reads of length 150 for ranibizumab-bevacizumab panning round 3 from 2 sample preparation each with 2 technical replicates, in which 82.9% were successfully parsed. The reduced sequence diversity in subsequent rounds indicated selection was occurring as expected.

To reduce noise in the training data for ranibizumab, we only retained sequences that had more than 5 read counts in at least one panning round, or had non-zero reads in all rounds. We then built 4 datasets for training machine learning models. In the classification dataset sequences with a round 3 frequency > 5e-5 or that is higher than its round 2 frequency by more than 1e-6 was labeled as positive, while sequence whose round 3 frequency is lower than round 2 frequency by more than 1e-6 and whose round 3 frequency is < 5e-5 was labeled as negative. This resulted in a training set of 51,130 sequences, with a distinct separation between the positive and negative examples. In the regression datasets, the R2-to-R3 enrichment was used as label and each dataset comprised subset of the whole pool. We added a pseudo-count of 1 to the read counts of all sequences to avoid errors when computing the ratio. The average size of the regression sets was 64,856.

For the multi-output training set, we retained sequences that had more than 3 read counts in at least one panning round and computed the R2-to-R3 enrichment as regression label for each of the targets. We then concatenated the labels, allowing missing values in one of the targets, and balanced the positive and negative samples by down-sampling negative/missing value sequences to have at most twice the size of the positive samples. This resulted in a training set of size 121,485 for etanercept and trastuzumab.

In the experiment with 1:100 diluted library of ML-proposed sequences to synthetic library sequences, we obtained 6,365,482 MiSeq reads with length 150 for standard washing condition and 7,016,792 reads for stringent washing condition for all 3 rounds. Over 85% of the reads were parsed successfully. We again observed decreasing diversity in subsequent panning rounds, and the most significant change of diversity happened between round 1 and round 2 where the number of unique sequences dropped from 1,044,401 to 76,928 for standard washing and from 1,135,356 to 53,809 in stringent washing. Our hypothesis was that the inclusion of highly competitive sequences generated by machine learning methods led to a more drastic competition in early rounds and thus accelerated the selection of good binders. Therefore, we looked at the R1-to-R3 enrichment in addition to R2-to-R3 enrichment when evaluating the enrichment for this experiment.

**Training an ensemble of neural networks**

We used six different architectures, five of which were convolutional neural networks with 1 or 2 convolutional layers with filter size of 1, 3 or 5 residues and stride 1, followed by local max-pooling layer with window size 2 and stride 2. We used 64 and 32 convolutional filters for single convolutional layer networks. In one of the double convolutional layer networks, we used 32 filters with width 5 in first layer and 64 filters with width 5 in the second layer. In the other network, we used 8 convolutional filters with width 1 in first layer to learn an embedding from one-hot to hidden space for each amino acid, and then used 64 filters with width 5 to learn higher level patterns. In each of the convolutional neural network, the output from last convolutional layer was fed into a fully connected layer with 16 hidden units and a dropout layer. It is then connected to the final output layer where the loss function is evaluated. We also designed a 2-layer fully connected neural network with 32 hidden units and dropout in each layer. Table S1 illustrates the detailed setup of each architecture and an estimation of the number of parameters in each architecture.

All the neural networks were implemented using Keras [27]. Training was conducted with respect to the binary cross entropy loss function in our classification task and the mean squared error loss in the regression tasks. For the multi-output regression networks, we used a masked mean squared error loss to deal with the missing values during training. Model hyperparameters were tuned by grid search using 10% of the training set as validation set and 5 epochs of quick training. The hyperparameters that yielded best validation loss were chosen for the training phase on the complete training set. We trained each neural network for 20 epochs using RMSProp optimization [28], and retained the learned network parameters from the epoch that produced best performance on the held-out test set.

**Optimizing antibody sequences with gradient ascent and a neural network ensemble**

Given that direct optimization on one-hot discrete space is difficult and inefficient, we used the gradients back-propagated from the neural network output layer to the input layer to guide the improvement of an input sequence. We relaxed the one-hot constraint during optimization to fully take advantage of gradient methods, periodically projecting the current continuous representation back to a discrete one-hot input by taking the arg-max of each amino acid position. By tracking the improvement and checking the convergence in one-hot space, we made sure that the optimization over one-hot space was happening along with the continuous space. By choosing different step sizes $\lambda$ and projection intervals k, we were able to control the level of divergence between proposed sequences and seed sequences. The gradient ascent approach can easily be applied to batches of seed sequences to fully utilize the parallel computing power of GPUs, and thus is efficient.

Because of the deterministic characteristic of gradient ascent, a single network can only produce limited number of sequences following the gradients determined by its local function landscape. We used an ensemble of 24 networks trained with different initialization and different subset of sequences, such that each of them learns a landscape with different "perspectives". This allowed us to optimize a single seed sequence with multiple potential directions and thus produced much more diverse candidates than a single network. We also varied the optimization hyperparameters, namely the step size ($\lambda$=1e-4, 5e-4, 1e-3, 5e-3, 1e-2) and projection intervals (k=10, 20, 30, 40, 50, 60, 70, 80), in order to produce diverse sequences of varying distances to their corresponding seed.  In total, we used 960 optimization methods to generate a pool of candidates.

A single neural network may overfit a training set given that a large portion of sequence space is not observed in our training data.  We used a two-step voting-thresholding strategy with an ensemble to increase the robustness of our optimization results to tackle this issue. First, we used all 960 optimization methods to vote on optimal sequences.  Each method voted for each sequence it produced, and the sequences with the top number of votes were selected for next step.  This voting step prioritizes sequences that were optimized with parameters that are neither too conservative nor too aggressive, and sequences that were constantly proposed by networks trained from different perspectives.  On the following step, we used an ensemble consisting of all 18 regression neural networks trained as before to evaluate each candidate.  Different networks tend to disagree in the regions where model uncertainty is high, and thus can be used to increase the robustness of enrichment prediction for unseen sequences.  We threshold the candidates by the 5% confidence lower bound of the scores obtained from the 18 neural nets, and chose the sequences with non-negative lower bound as the positive candidates that will be used for comparison with the seeds.  We used sequences with negative scores as negative controls to validate the performance of computational models.

**Stringent washing improves antibody EC50 prediction**

The enrichment of a CDR-H3 sequence in a phage panning experiment is only an approximation of affinity for a target.  During panning Fab fragments displayed on phages specific for the target are competing with both non-binding and poly-specific (“sticky”) Fab fragments.  Standard washing conditions are designed to reduce non-binding Fab fragments with multiple washing cycles.  Poly-specific Fab fragments are depleted by using blocking agents (Supplemental Methods).

We hypothesized that we could improve the correspondence between phage panning enrichment and IgG affinity by using stringent washing condition during phage panning to reduce non-specific interactions.  We defined a stringent wash buffer with increased ionic strength to interrupt salt bridges (0.75M NaCl) and with a detergent to reduce hydrophobic interactions (2% Octyl β-D-glucopyranoside) in 0.1M Tris at pH 7.4.  We expected that this stringent washing buffer would reduce both non-specific binding and weak specific binding. Weak specific binding can result from a low number of ionic and / or hydrophobic target interactions.

We performed panning experiments against ranibizumab using this stringent wash buffer starting in the second round of panning. Selected sequences at differing enrichments were produced as full-length IgG molecules, and used in an ELISA EC50 [29] assay as a measure of antibody affinity. We found that the stringent buffer increased the correspondence between panning enrichment and affinity (Table S3, Fig. S3A). The Spearman rank correlations between enrichment and affinity for the sequences in Table S3 are 0.266 ($p = 0.404$, $n = 12$) for standard washing condition, and 0.371 ($p = 0.236$, $n = 12$) for stringent washing condition.

**Alternative machine learning methods for proposing sequences**

In addition to our neural network ensemble, three other machine learning frameworks were also applied to generate novel CDR-H3 sequences. Here, we detail these alternative computational approaches to antibody design that were less successful than our Ens-Grad methodology.

Genetic algorithms for antibody sequence optimization

As an alternative approach to our Ens-Grad methodology, we also applied a non-gradient-based optimization procedure to propose high affinity CDR-H3 sequences. We designed two genetic algorithms (GA-KNN and GA-CNN) that seek to explore sequences under the guidance of a fitness function that estimates the enrichment of a proposed sequence. The two genetic algorithms differed only in their fitness functions that were used to estimate the predicted enrichment of a CDR-H3 sequence.

Each genetic algorithm selected random sequences to start. In each generation, the genetic algorithm mutated parental sequences 600 times and performed a crossover on a pair of parents 600 times to produce a total of 1200 new children. In practice, we observe that random crossovers and mutations are quite unlikely to produce sequences that resemble valid antibody sequences in the seed pool. To enhance the sampling efficiency of the genetic algorithm, random crossover is then performed between each child sequence generated and its 10 nearest neighbor sequences in 1000 randomly sampled sequences from the seed pool. This effectively "projects" back the sampled child sequence back to the space of allowable sequences. After each child sequence is processed in this way, 3000 survivors from the 4200 sequences were probabilistically chosen based on fitness. Two fitness functions were evaluated. For GA-KNN we used the average scores of the ten nearest neighbors from 1000 examples in the training set. For GA-CNN, we used the prediction from a single convolutional neural network that was trained on the training set and consists of a convolutional layer of 64 filters with size 5 and stride 1, a max-pooling layer of size 2, a fully-connected layer of 16 neurons, and a Dropout layer with a dropout rate of 0.3 . The genetic algorithms ran for 30 generations. The final pool of generated sequences consists of all the sequences the algorithm sampled.

Variational autoencoder model for antibody sequence optimization

Another alternative approach to Ens-Grad we also applied for optimizing antibodies was a framework based around a recurrent variational autoencoder (VAE) generative model that has been proposed to revise natural language and other discrete sequence structures [30]. This approach is based around the idea that combinatorial optimization of discrete sequences may be easier in an alternate space of learned continuous representations, in which both the fitness function and the region of high-performing sequences adhere to a smoother and simpler geometry. Before optimization of antibody sequences, the VAE approach first learns a probabilistic generative model of the antibodies, in which continuous latent vectors are responsible for producing both the antibody CDR-H3 sequence as well as the corresponding binding affinity. Both the likelihood function of sequences given latent factors (decoder) as well as an inference model of latent values responsible for generating a particular sequence (encoder) are parameterized using GRU recurrent neural networks [31], which are jointly trained with a feedforward neural network that predicts binding affinity from inferred-values of the latent factors (Fig. S4). After these components are learned, we leverage this model to optimize one of the seed CDR-H3 sequences as follows: the sequence is encoded into the latent space by our inference network, then constrained gradient-ascent is applied to these continuous latent variable values with respect to the predicted binding affinity until a locally optimal latent-factor configuration is found. Finally, the most likely sequence (according to our decoder network) associated with this optimal latent vector is returned as the optimized sequence. This method simultaneously leverages the generative capacities of the recurrent decoder (to ensure natural-looking CDR-H3 sequences are produced), the compressed sequence representation learned by the recurrent encoder network (to enable the application of lower-dimensional continuous optimization algorithms), and the posterior Kullback-Leibler penalty of the VAE (to ensure the encoded sentence representations follow a simple geometry).

Our VAE method was applied with the same architecture, training, and sequence-optimization settings detailed in [30]. We employed 128-dimensional vectors as latent factors together with a learned 8-dimensional continuous representation of each amino acid upon which our inference network was trained to operate. Training proceeded with the mean-squared error loss applied to binding affinity labels equal to the R2-to-R3 enrichment, as in the previously described regression task. From each seed sequence, this method was applied to produce at most

20 new sequences, where generated sequences that violated our set of structural conditions were subsequently omitted as in the case of the other methods.

#### Enforcing compositional constraints in sequences used for oligo synthesis

After combining the optimized sequences proposed by all synthetic methods and the control sequences, we applied a post-processing pipeline to all sequences before oligo synthesis. The post-processing pipeline removes sequences with cysteine and filters out sequences with Asn-Gly, Asn-Ser, Asn-X-Ser or Asn-X-Thr for reasons described in the library construction section. In order to produce valid CDR-H3 sequences that can be folded correctly, we constrained the last two positions in the CDR-H3 sequences to be fixed as the framework suggested. We also avoided proposal of cysteine or asparagine during ensemble gradient optimization and VAE optimization, due to the aforementioned reasons.

#### Oligo synthesis of CDR-H3 sequences and library generation

We constructed a library including 9,553 sequences proposed by ensemble gradient ascent (Ens-Grad). 5,467 of these sequences were optimized from denoised seeds and thus were included in the analysis to avoid over-estimation of the performance. We also included 9,503 sequences proposed by VAE, 17,813 sequences proposed by GA-KNN, 6,286 sequences proposed by GA-CNN, and 23,606 sequences designed for other purposes and thus not described in this paper, summing up to 64,984 machine learning derived sequences. Here the number is the total number of sequences proposed by each method even though there were overlapping sequences between different methods that are only counted once when calculating the library size. We also included 39,541 control sequences consisting of 12,243 seed sequences defined as previously described and 27,298 controls which contained both negatively-enriched sequences from our panning experiment (negative control) as well as additional sequences generated to receive negative predictions from our neural network ensemble. Collectively we built a library of 104,525 CDR-H3 variable region sequences. We concatenate the sequences with the framework and restriction sites used for cloning the sequences into phages. We translated the amino acid sequences into DNA nucleotide sequence using EMBOSS [26] and the E.coli codon usage table was used. The nucleotide sequences were then used to synthesize a DNA oligo pool (Twist Bioscience). Library inserts were generated by PCR using Phusion High Fidelity DNA polymerase (NEB Biolabs). The resulting CDR-H3 library inserts were ligated into the base template and transformed into E.coli TG1 DUO (Lucigen) aiming for a library size of 5E+08 transformants. Phages were produced using VSCM13 helper phage (Agilent) using standard previously described protocols [23].

**Stringent de-noising of seed sequences**

To provide a conservative estimate of the improvement caused by machine learning we used an extra step of noise correction to ensure that our measures of machine learning performance were not confounded with beneficial random noise caused by sequencing errors. We eliminated seeds for optimization that contained sequencing errors by coalescing clusters of reads that were predicted to be derived from a single parent sequence.

The models we used for optimization were trained on data that did not have sequencing errors removed so as to not advantage their performance. We found that model performance when trained on de-noised data was comparable to model performance when trained on data containing sequencing errors (Fig. S5).

Assuming the PCR and sequencing error rate per base is $e$, the probability of getting a particular $k$-error read from a $L$-nt sequence is:

$$p_{Lk} = e^k(1-e)^{L-k}$$

Given $N$ reads of a $L$-nt sequence, the occurrence of a particular $k$-error sequence follows a Binomial distribution with parameters $(N, p)$. Thus the probability that the read count of a particular $k$-error sequence is less than $M$ is

$$q_{NMLk} = \sum_{m=0}^{M-1} \binom{N}{m} (p_{Lk})^m (1-p_{Lk})^{N-m}$$

The $k$ errors can locate in $\binom{L}{k}$ different places. Assuming $M \ll N$, we can consider the read counts of different k-error sequences as identically and independently distributed variables. Then the probability of observing at least one $k$-error sequence with $M$ or more reads can be approximated as:

$$P_{NMLk} = 1 - (q_{NMLk})^{\binom{L}{k}}$$

For each sequence, we identified the most abundant sequence that is $k$-bp away (for $k \in \{1,2,3\}$) and consider them as potential parent sequences. For each potential parent-child sequence pair, we calculate $P_{NMLk}$ with $N$ being the sum of the read counts from the parent and child and $M$ being the read count from the child. Assuming the number of sequences to de-noise is $S$, we performed FDR correction (Benjamini–Hochberg) to identify pairs with FDR>0.05. Any child with such pairs are considered noisy reads generated from the corresponding parent.

Using a literature [32] suggested per-base error rate of 0.001, we performed the above de-noising procedure on reads combined from round 2 and round 3 of the initial panning experiment for a higher signal-to-noise ratio. The DNA sequences that survived the de-noising procedure and don't contain uncertain bases (N) were translated and intersected with the seed sequences to obtain the final set of de-noised seeds.

**Visualization of sequences using Parametric t-SNE**

t-Distributed Stochastic Neighbor Embedding (t-SNE) [33] is a widely-used technique for dimension reduction that is particularly well suited for visualizing high-dimensional datasets. Specifically, given a set of $D$ dimensional vectors $X_1, X_2, \ldots, X_n$, for any pair of points $X_i$ and $X_j$, t-SNE computes a conditional probability $p_{i|j}$ that $X_i$ would pick $X_j$ as its neighbor if the neighbors were picked in proportion to their probability density under a Gaussian centered at $X_i$:

$$p_{i|j} = \frac{e^{-\frac{dist(x_i,x_j)^2}{2\sigma_i^2}}}{\sum_{k\neq i} e^{-\frac{dist(x_i,x_k)^2}{2\sigma_i^2}}}$$

Then t-SNE computes a similarity matrix $P$ where the similarity between any pair of $X_i$ and $X_j$ is defined as:

$$P_{ij} = \frac{p_{i|j} + p_{j|i}}{2}$$

Given the similarity matrix, t-SNE aims to embed each $X_i$ into a d-dimensional vector $Y_i$ such that the similarity between any pair of points is maintained as well as possible. For a given set of d-dimensional embedding $Y_1, Y_2, \ldots Y_n$, t-SNE considers the distance between any pair of $Y_i$ and $Y_j$ follows a Student t-Distribution with one-degree of freedom, and the similarity between $Y_i$ and $Y_j$ is proportional to the probability density of the distance:

$$q_{ij} = \frac{(1 + dist(y_i, y_j)^2)^{-1}}{\sum_{k\neq i}(1 + dist(y_i, y_j)^2)^{-1}}$$

t-SNE learns a set of embedding $Y_1, Y_2, \ldots Y_n$ that maintains the pairwise similarities by minimizing the Kullback-Leibler divergence between the two similarity matrices p and q:

$$KL(p||q) = \sum_{ij} p_{ij}\, log \frac{p_{ij}}{q_{ij}}$$

To visualize the CDR-H3 sequences, we used t-SNE to embed each sequence into a 2-D vector. We used edit distance as the distance metric for the original sequence space ($X_1, X_2, \dots, X_n$), and euclidean distance for the embedded 2D vector space ($Y_1, Y_2, \dots Y_n$).

One major challenge of t-SNE is the lack of scalability. It is infeasible in time and memory to apply t-SNE on hundreds of thousands of points as we have in our dataset. Therefore, we instead learn a fixed embedding function that maps a given input $X_i$ to the corresponding embedding $Y_i$. We parametrized this function using a Residual Network (ResNet) [34], and trained the network by minimizing the same Kullback-Leibler divergence objective as t-SNE. A similar idea that uses less expressive network architecture and more complicated training procedure has been explored in the computer vision community with promising results [35].

Compared to t-SNE, we are now able to perform visualization on datasets with a large number of samples thanks to three run-time improvements. First, with highly expressive neural network structure such as ResNet, the number of parameters we learn is sub-linear in the number of training samples. Second, with optimization algorithms that are based on stochastic gradient descent (SGD), the neural network can be trained in a memory-efficient way by only observing a "mini-batch" of the dataset in each iteration. Third, parameter optimization can be performed efficiently using Graphics Processing Units (GPUs). We note that unlike the conventional training of neural networks, a large mini-batch size is needed to effectively approximate the distribution of the similarities in the whole dataset. In this study, we used a ResNet with one convolutional layer followed by four residual blocks and a 32-neuron fully connected layer before the final output layer. Each residual block predicts the residual between the output and the input with three sets of convolutional, batch-normalization, and ReLU-activation layers. All the convolutional layers in this model have 64 filters of width 3. Training was performed with a batch-size of 6000 using Adam, an adaptive stochastic gradient method [36].

We note that more recent visualization techniques, such as UMAP[37], also have the potential to scale to the hundreds of thousands of samples that we have. Compared to the alternative techniques, parametric t-SNE conveniently learns a fixed mapping from a CDR3 sequence to the corresponding low-dimensional representation. Such a mapping allows us to apply the same transformation to CDR3 sequences from different panning rounds to elucidate how the sequence space evolves as the panning experiment progresses.

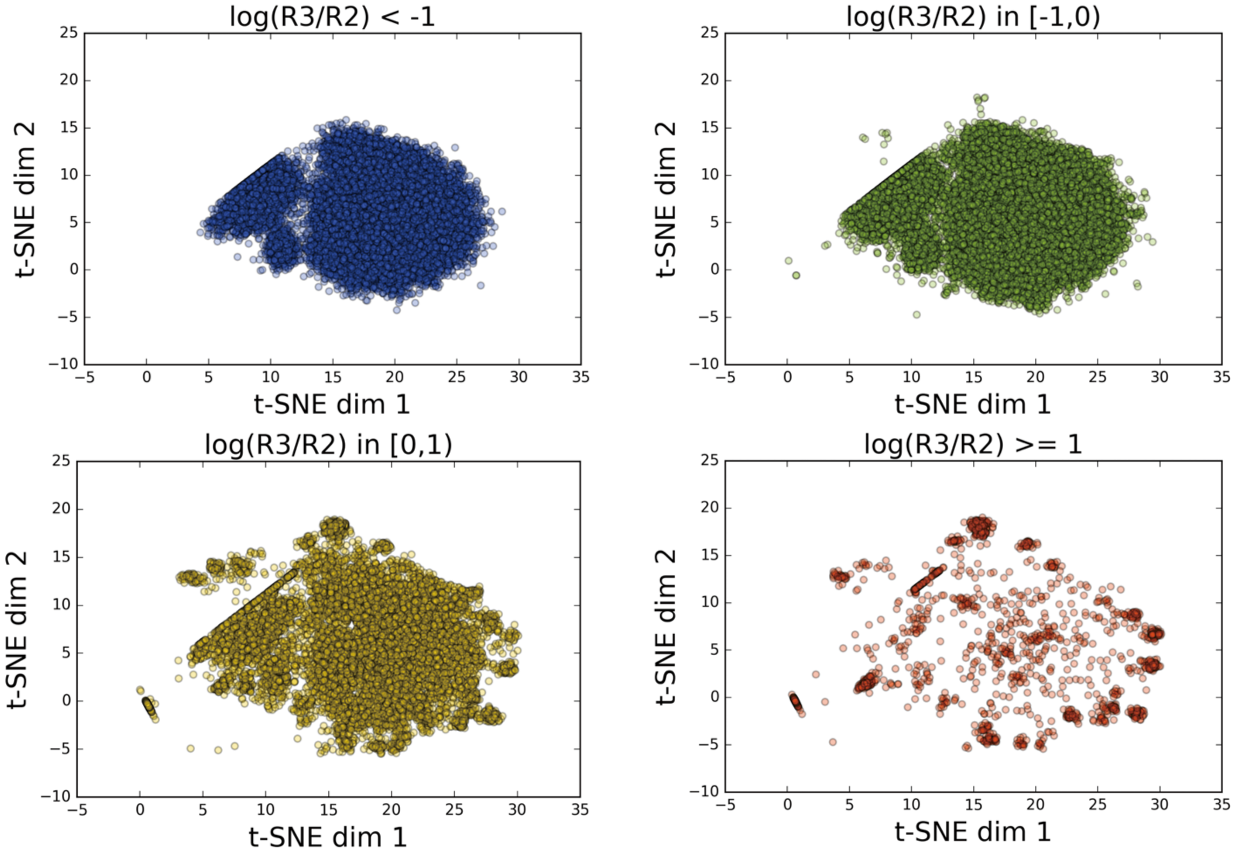


**Fig. S1.** t-SNE visualization of all valid sequences in round 2 and round 3 in initial synthetic library grouped by their enrichment. High enrichment sequences tend to have more clustered structures compared to non-binders.

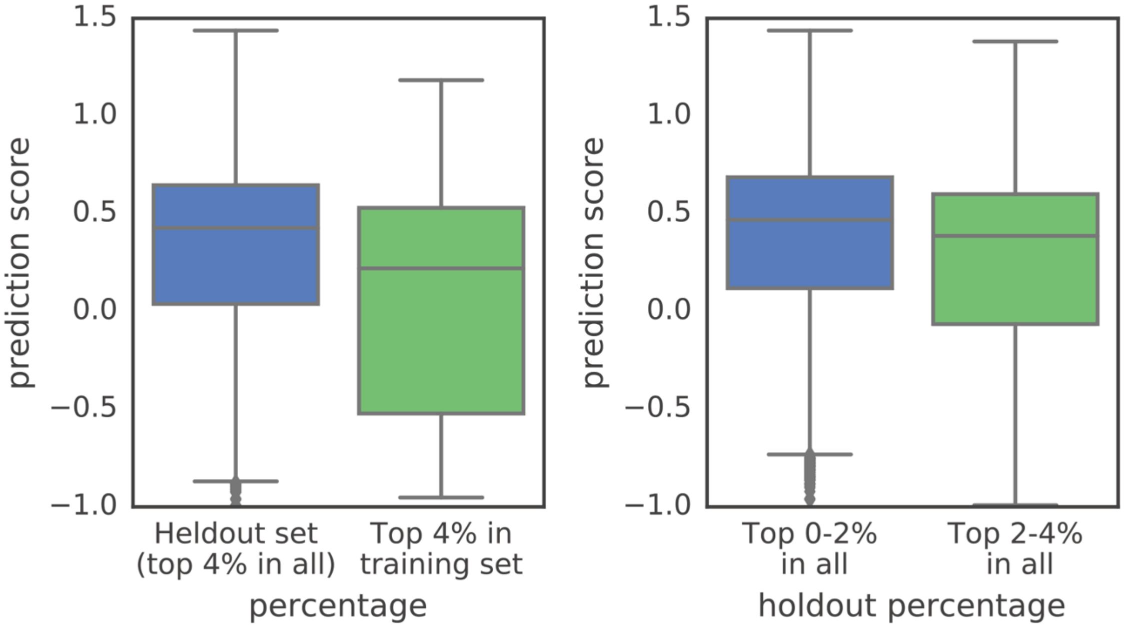


**Fig. S2.** Convolutional neural networks can accurately extrapolate to new sequences which are more positively enriched than the entirety of the training set. We held out the top 4% of the sequences and trained the model solely using the bottom 96% of sequences with the lowest enrichment values. Our convolutional network assigns higher prediction scores to the held-out top 4% of all sequences than to sequences in the top 4% of its reduced training set (two-sided Mann–Whitney U test, $U = 4485500$, $p < 2.2e-16$, and difference in location (Hodges–Lehmann estimate) = 0.172 with 95% confidence interval = [0.147, 0.198]). The model is also able to differentiate between sequences whose enrichment lies in the upper vs. lower half of this held-out top 4% set (two-sided Mann–Whitney U test, $U = 1049600$, $p = 1.036e-10$, and difference in location = 0.0974 with 95% confidence interval = [0.0676, 0.127]). Boxplots show median (the center line in the box), 25th and 75th percentiles (the boundaries of the box), 1.5 interquartile range (the ends of the whiskers), and outliers (points outside of the whiskers).

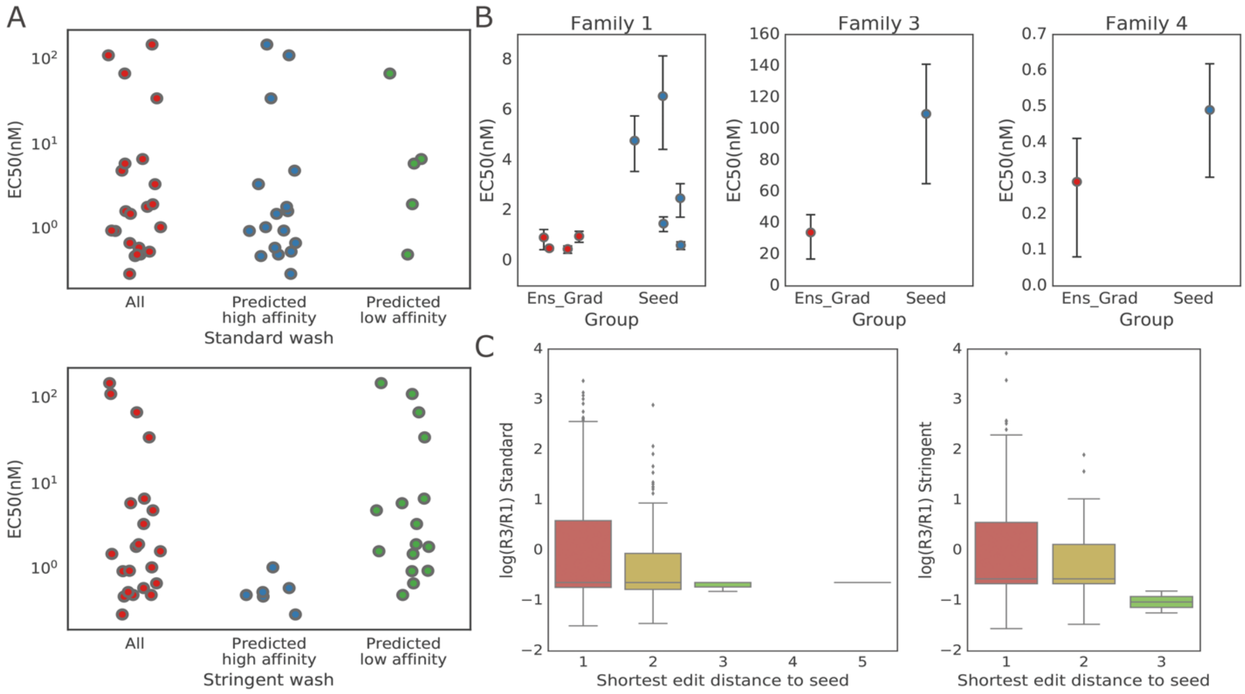


**Fig. S3.** (A) EC50 correlation with standard (upper) and stringent (bottom) washing condition. The stringent washing condition provides enrichment values that are better aligned with actual EC50 measurements, whereas standard washing has more false positives. (B) EC 50 of subsets of top Ens-Grad proposed sequences and seed sequences in four different families with 95% confidence error bars (the whiskers in the plot). (C) Distribution of log10(R3/R1) enrichments of Ens-Grad proposed sequences with different shortest edit distances to a seed sequence. The distribution of R1-to-R3 enrichment is shown vs. edit distance for (A) standard washing condition, and (B) stringent washing condition. Boxplots show median (the center line in the box), 25th and 75th percentiles (the boundaries of the box), 1.5 interquartile range (the ends of the whiskers), and outliers (points outside of the whiskers).

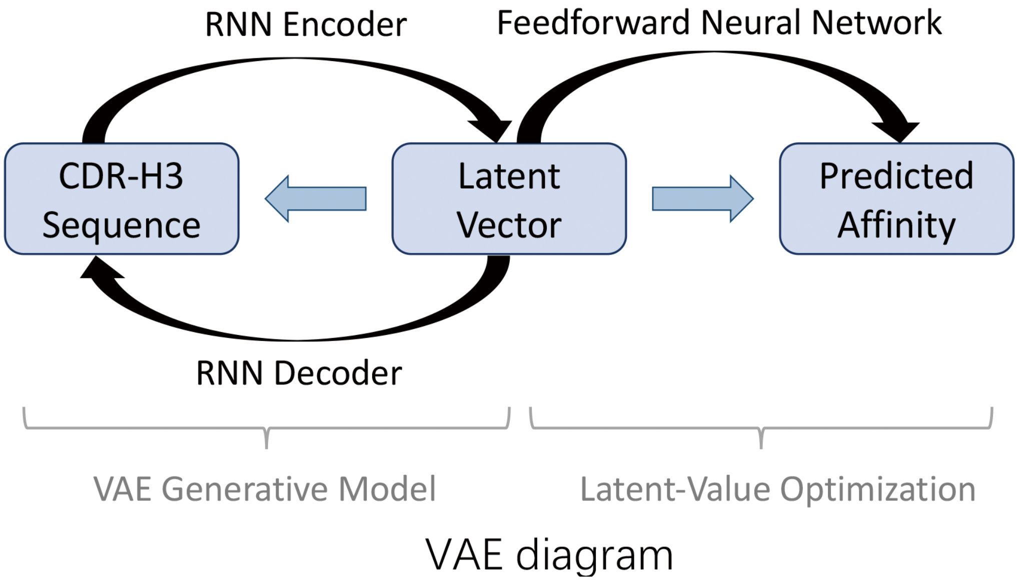


**Fig. S4.** Overview of the alternative VAE approach to produce high-affinity sequences. Sequences are encoded in a continuous latent representation that is used to predict affinity via a simple output function, and the optimization to identify higher-affinity sequences is performed over these latent vectors.

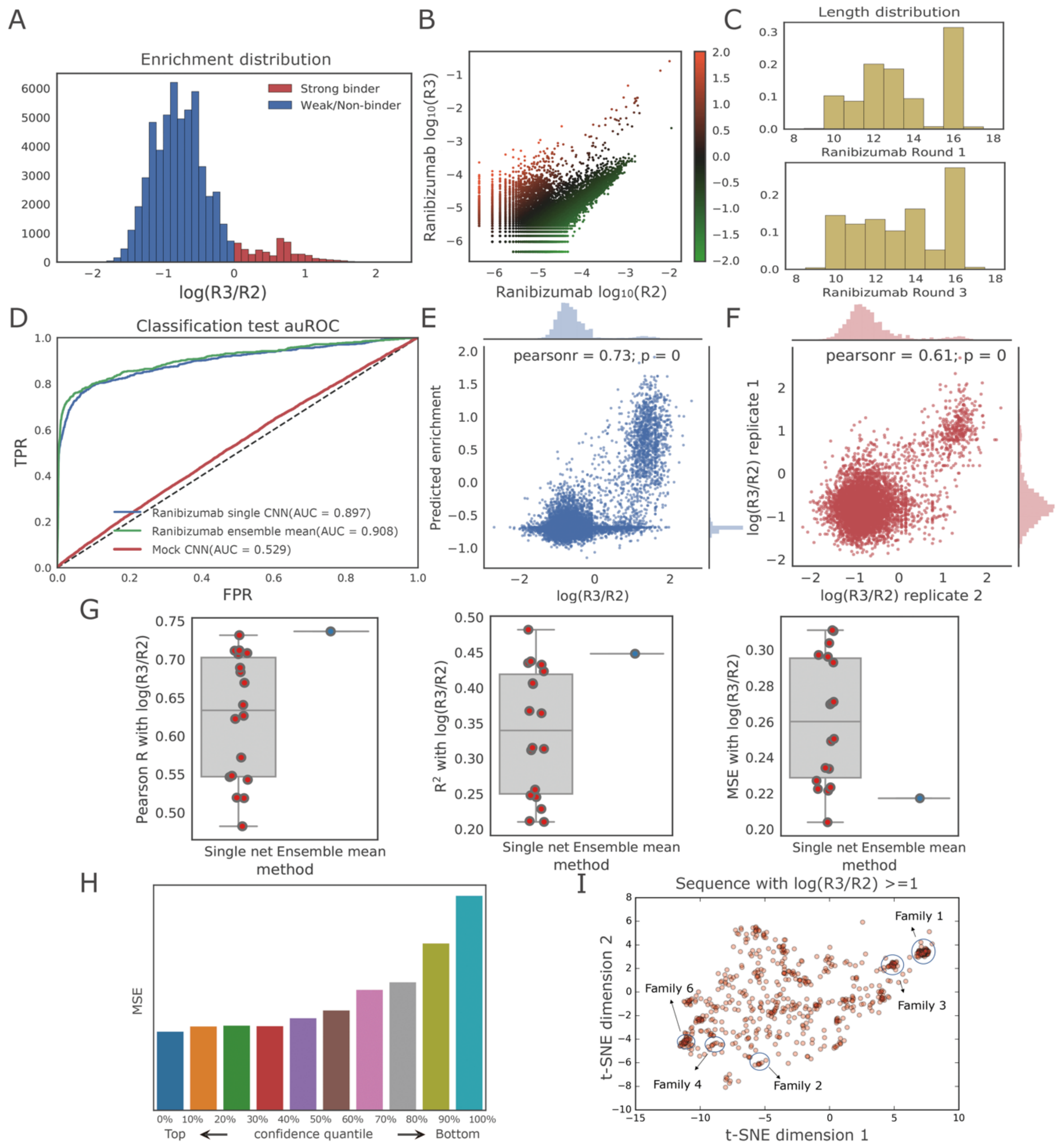


**Fig. S5.** Summary of the training data processed with an extra step of de-noising and prediction performance of our neural networks trained on denoised ranibizumab data. (A) Histogram of R2-to-R3 enrichment in training data. R2-to-R3 enrichment is defined as log10 of R2-to-R3 frequency ratio. (B) Scatter plot of log10 sequence frequency in R2 and R3, colored by the R2-to-R3 enrichment value. Each point represents a unique valid sequence in the NGS output, where points above the diagonal have positive R2-to-R3 enrichment and vice versa. (C) Histogram of CDR-H3 sequence length before and after 2 rounds of panning. (D) ROC curve of classification task using the best single network and the ensemble, compared with a mock control. The ensemble method outperforms a single neural network, while the performance in mock is close to random guessing, as expected. (E) Regression performance on non-overlapping sequences in replicate II using the best single network (Pearson $r = 0.73$, $p < 1e\text{-}16$, $n = 19{,}546$). (F) Replicate consistency of enrichment (Pearson $r = 0.61$, $p < 1e\text{-}16$, $n = 5{,}677$). (G) Ensembles produce estimates of enrichments that are better than any single network in terms of Pearson r and competitive in terms of R2, and mean square error. Boxplots show median (the center line in the box), 25th and 75th percentiles (the boundaries of the box), 1.5 interquartile range (the ends of the whiskers), and outliers (points outside of the whiskers). (H) Uncertainty estimation using ensemble correlates with the prediction accuracy. Sequences with higher confidence have lower mean square error. (I) t-SNE visualization of sequences with positive enrichment between round 2 and round 3. The families in Table 3 are labeled by circles. CDR-H3 sequences with high enrichment form isolated clusters. binders (survived sequences in round 3) showed preferred CDR-H3 lengths when comparing to sequences in round 1.

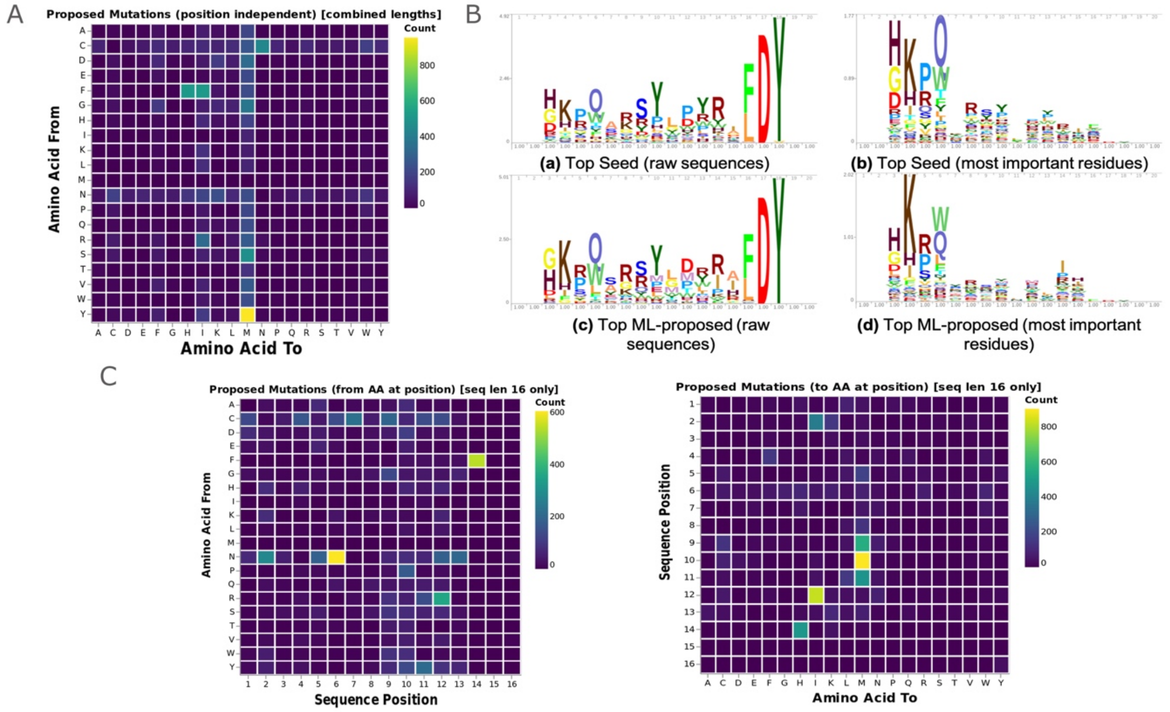


**Fig. S6.** Visualization of machine learning (ML) optimization. (A) Position independent proposed mutations map. Heatmap visualization showing the count of observed sequence mutations between seed and corresponding ML-proposed sequences. The mutation count indicates the number of mutations observed from/to each amino acid in 7614 pairs of seed and ML-proposed sequences, regardless of the position of the mutated residue. Cysteine and asparagine are not proposed by our ML method (Methods) (B) Sequence logo visualizations for top 1000 sequences of length 16 based on mean predicted R3/R1 enrichment from the neural network ensemble. Sequence logos for Seed (a) and ML-proposed (c) groups are based on residue frequency, and sequence logos Seed (b) and ML-proposed (d) are based on frequency of residues marked as most important for predicted sequence enrichment. Sequence logos are computed using Skylign. (C) Position dependent proposed mutations map (sequences length 16 only). Heatmap visualization showing the count of ML-proposed sequence mutations occurring from (left) and to (right) each amino acid at each sequence position. The mutation count indicates the number of mutations observed from/to each amino acid at each sequence position in 5025 pairs of seed and ML-proposed sequences having length 16.

| | Num. of convolutional layers | Conv 1 | Conv 2 | Num. of fully connected layer | Num. of Fully connected neurons | Total Num. of parameters |
|---|---|---|---|---|---|---|
| **Seq_32_32** | **0** | **N\A** | **N\A** | **2** | **32** | **13954** |
| **Seq_32x1_16** | **1** | **Width 5, 32** | **N\A** | **1** | **16** | **8402** |
| **Seq_32x2_16** | **2** | **Width 5, 32** | **Width 5, 64** | **1** | **16** | **18706** |
| **Seq_64x1_16** | **1** | **Width 5, 64** | **N\A** | **1** | **16** | **16754** |
| **Seq_32x1_16_filt3** | **1** | **Width 3, 32** | **N\A** | **1** | **16** | **7122** |
| **Seq_embed_32x1_16** | **2** | **Width 1, 8** | **Width 5, 32** | **1** | **16** | **13082** |

**Table S1.** Neural network architectures used in the Ens-Grad model ensemble and the number of parameters in each neural network model.

| | before | after |
|---|---|---|
| **Total bevacizumab binders** | 27432 | 22037 |
| **Ground truth Fc binders** | 366 | 26 |
| **Non-specific binders** | 1530 | 380 |
| **Specific bevacizumab binders** | 25902 | 21657 |

**Table S2.** Number of ground truth bevacizumab Fc binders and non-Fc binders before and after applying Fc filter using ensemble neural networks on all panning derived bevacizumab binders. The total number of panning derived bevacizumab binders is also included. We observed that the computational filter eliminated significantly more non-specific binders than specific binders, suggesting that machine learning models can target non-specific binding precisely.

| CDR-H3 Sequence | Group | $R^2$ | EC50(nM) | Standard log(R3/R1) | Stringent log(R3/R1) |
|---|---|---|---|---|---|
| **VRGGHEFEKRVHDY** | Primary | 0.9919 | 2.59 | 2.356 | 3.016 |
| **GHYYDIGVFPWDTFDY** | Primary | 0.9897 | 0.49 | 2.467 | 2.901 |
| **HKPQAKSYLPYRILDY** | Primary | 0.9917 | 6.55 | 3.068 | 2.703 |
| **GKSLYGQETTWPHFDY** | Primary | 0.9901 | 0.67 | 3.241 | 2.671 |
| **WQQWSGYPRQKYSFDY** | Primary | 0.9753 | 66.81 | 2.4 | 2.397 |
| **GVHYYWSYPRSATFDY** | Primary | 0.9224 | 15.11 | 1.658 | 2.251 |
| **YRSPHHRGGATWQFDY** | Primary | 0.9919 | 5.79 | 3.067 | 1.843 |
| **QQAQVISYYSYSTFDY** | Primary | 0.9945 | 16.01 | 2.119 | 1.618 |
| **GIEGAWYSYKPHKFDY** | Primary | 0.9621 | 16.44 | 1.823 | 0.934 |
| **GKRWSRRYGDRRAFDY** | Primary | 0.9895 | 1.47 | 3.293 | 0.418 |
| **LHRWSRRWGWDRRFDY** | Primary | 0.9915 | 1.12 | 3.418 | 0.361 |
| **DLFRYYYFFWPLDY** | Primary | 0.9896 | 109.5 | 3.433 | 0.28 |

**Table S3.** Examples of the top primary sequences derived by the initial phage panning using synthetic library along with their enrichment in standard and stringent washing condition, corresponding mean EC50 affinity measurement, and R2 of the replicate EC50 measurements. The Spearman rank correlations between enrichment and EC50 are 0.27 (standard wash) and 0.37 (stringent wash).
